# Protection from plague via single dose administration of antibody to neutralize the type I interferon response

**DOI:** 10.1101/2024.03.11.584497

**Authors:** KD Marks, DM Anderson

## Abstract

*Yersinia pestis* is a gram-negative bacterium and the causative agent for the plague. *Yersinia spp*. use effector proteins of the type III secretion system (T3SS) to skew the host immune response toward a bacterial advantage during infection. Previous work established that mice which lack the type I IFN receptor (IFNAR), exhibit resistance to pulmonary infection by *Y. pestis*. In this work, we addressed the efficacy of a single dose administration of neutralizing antibody to IFNAR (MAR1) as a preventive treatment for plague. We show that single dose administration of MAR1 provides protection from mortality due to secondary septicemic plague where it appears to reduce the production of serum TNFα during the disease phase. We further demonstrate that the T3SS effector protein YopJ is necessary for MAR1-induced protection, however IFNAR-dependent serum TNFα was observed independent of YopJ. We further define tissue-specific anti-bacterial roles of IFNAR that are blocked by YopJ activity indicating that YopJ and IFNAR work in parallel to promote disease. The combined data suggest that therapeutic targeting of IFNAR signaling may reduce the hyper-inflammatory response associated with plague.

## Introduction

*Yersinia pestis* is a gram-negative facultative intracellular pathogen and the causative agent for the plague [1]. Bubonic plague is the most common form of the disease, transmitted to humans via the bite of an infected flea. Pneumonic plague occurs following inhalation of *Y. pestis*, is characterized as a complete disarmament of the innate immune response and logarthimic growth of *Y. pestis* [2, 3]. Systemic spread of *Y. pestis* through the vasculature is often a consequence of bubonic or pneumonic plague, with the development of severe sepsis commonly observed as the cause of death. All forms of the disease involve necrosis of immune tissues, inflammatory toxicity, and an acute phase response. Consequently, the disease often continues even when antibiotic treatment successfully resolves infection [4].

Although most naturally occurring *Y. pestis* strains are sensitive to multiple classes of antibiotics, the treatment of plague by antibiotics is often unsuccessful. Therefore the global mortality rate of plague in the 21^st^ century is 10-20%, with some recent outbreaks having mortality rates as high as 75% [5-7]. While it is clear that early antibiotic treatment is associated with improved outcome, the rise of naturally occurring antibiotic resistance is concerning [8]. Isolates from Madagascar and China have been recovered from patients in the late 20^th^ and early 21^st^ centuries that exhibited multi-drug resistance. Mechanisms of resistance include mutations as well as horizontally acquired genes [9-14]. These naturally evolved strains or the genetic engineering of multi-drug resistant *Y. pestis* could lead to the spread of plague with high mortality rates. Therefore it is necessary to continue to prioritize research towards understanding the pathogenesis, prevention and treatment of plague.

Plague is characterized as a rapidly progressing disease with an extremely low infectious dose, two features that depend on the regulated expression of the type III secretion system (T3SS) which is encoded by the pCD1 plasmid [15]. This virulence factor is utilized by extracellular bacteria and is engaged when *Y. pestis* makes intimate contact with a host phagocytic cell [16]. The T3SS is a conduit that spans across the bacterial cell wall and outer membrane and into host cells through which effector proteins (collectively known as *Yersinia* outer proteins, Yops) are translocated into eukaryotic target cells [17]. At least 7 Yops are known to have effector function in host cells, where they impact signaling pathways that otherwise operate to activate the innate immune response. These include the inhibition of phagocytosis, inflammatory cytokine production, and the inflammasome, while also sensitizing the immune cells to undergo programmed cell death [18]. The consequence of the T3SS effector proteins on the infection is an early anti-inflammatory effect that is permissive for extracellular bacterial growth [19].

In addition to being a part of the *Yersinia* family of effectors, YopJ and its homologs are found in the T3SSs of pathogenic bacteria, whether they infect animals or plants [20]. YopJ has been characterized as an acetyltransferase that becomes activated upon binding to inositol hexakisphosphate produced in host cells [21]. Transfer of CH_3_CO-from acetyl coenzyme A (AcCoA) to target proteins in the mitogen activating protein (MAP) kinase family results in their inactivation [22]. As these proteins are typically involved in stimulating inflammation, YopJ activity strongly suppresses production of pro-inflammatory cytokines. YopJ may also have deubiquitinase activity that contributes to the inactivation of MAP kinase signaling [23]. YopJ acetylation and inactivation of transforming growth factor β (TGFβ) activating kinase 1 (TAK1) causes the activation of non-canonical pyroptosis through caspase 8 [24, 25]. In contrast, accumulation of YopJ shifts the function of caspase 8, and results in the inactivation of the surface receptor for tumor necrosis factor α (TNFα) [26]. Despite these functions in cell death and inflammation *in vitro*, the importance of YopJ to the progression of plague is unclear. Deletion of *yopJ* from *Y. pestis* caused only a minor attenuation of virulence in the rat bubonic plague model, yet YopJ is found in all strains of *Y. pestis* indicating it is under positive evolutionary selection [27]. In contrast, hyper-injection of YopJ more severely attenuates *Y. pestis* virulence [28]. These observations suggest there may be opposing effects of YopJ activity on the host response to infection.

Amongst the myriad of host cell responses, the type I interferon (IFN) response is one of the more cogent methods of defense for innate and adaptive immunity. Initially characterized during viral infection, type I IFNs are produced and secreted as a consequence of infection or cancer. Virtually all cells are capable of producing type I IFN, and conversely virtually all cells express the type I IFN receptor, called the IFN αβ receptor (IFNAR) whether as an infected or bystander cell. The type I IFN response involves disruption of translation to prevent viral replication in the cytosol [29]. Type I IFN signaling also sensitizes infected or damaged cells to undergo programmed cell death. The transcriptome of the type I IFN response has been studied in numerous systems, and the total number of interferon stimulated genes (ISGs) varies depending on the model that was assayed. Cross-comparison of the data in the Interferome database suggests a core set of 69 ISGs, however as much as 10% of the genome may be influenced by the type I IFN response [30, 31].

Type I IFNs regulate the recruitment or activity of granulocytes during infection through numerous mechanisms, including IFN-dependent chemokines and cytokines that direct or suppress neutrophil recruitment [32, 33]. Neutrophils are the predominant immune cell type that contributes to eradication of *Y. pestis* and other bacteria, but excessive recruitment or activation of neutrophils is associated with disease [34]. During cancer, type I IFN signaling regulates activation, migration and lifespan of neutrophils in a beneficial manner towards natural antitumor immunity [35]. Global transcriptional analysis has demonstrated tissue specific and systemic roles of type I IFN in multiple immune cell subtypes that influence neutrophil recruitment and function in the lungs during bacterial infection [36, 37]. During *M. tuberculosis* infection, however, type I IFN has clear association with pathology. Patients with active tuberculosis show a type I IFN transcriptome signature, with severity of disease correlating with ISG over-expression in neutrophils and monocytes [38]. The degree to which this occurs during other infections is unknown.

We previously showed that the type I IFN response contributes to the pathogenesis of plague, with *Ifnar*^*-/-*^ mice exhibiting greater sensitivity to infection by *Y. pestis* KIMD27 [39]. We further demonstrated type I IFN expression in *Y. pestis*-infected lungs as well as in the serum of infected mice that can be detected in the early stage of infection as well as in the disease phase [40]. These data are consistent with multiple roles for type I IFN signaling throughout the course of infection. Furthermore, we found that the T3SS effector protein YopJ modulates IFNβ production in macrophages *in vitro*. Since IFNβ is an ISG, this result suggests that YopJ may be able to perturb type I IFN signaling. To distinguish these possibilities, in this work, we evaluated therapeutic inhibition of type I IFN signaling and the role of YopJ using a murine model of secondary septicemic plague.

## Methods

### Bacterial Strains and Load Quantification

*Yersinia pestis* KIM6-pCD1^Ap^ (referred to as KIM5-) is a non-pigmented (*pgm*-) mutant that is exempt from select agent registration [41]. The isogenic *yopJ*_*C172A*_ mutant strain carries a point mutation in YopJ that abrogates its function giving a null phenotype [42]. For generation of seed stock, *Y. pestis* strains were grown fresh from a frozen stock by streaking for isolation onto heart infusion agar (HIA) plates; single colonies were used to inoculate 40 mL of heart-infusion broth (HIB) with 2.5M of CaCl_2_, then grown at 37°C for 24 hours with aeration at 125 rpm. Multiple seed stocks were frozen in -80°C at OD_600_ of 0.2. For each experiment, seed stock was thawed and grown in the HIB liquid broth for 24 hours. For enumeration of *Y. pestis* in mice, tissues were homogenized in 1mL of sterile PBS, then serially diluted and plated in duplicate on HIA.

### Animal studies

**Ethics Statement:** All animal procedures were performed in compliance with guidelines of the Office of Laboratory Animal Welfare and the National Institutes of Health Guide for the Care and Use of Laboratory Animals and were approved by the University of Missouri Animal Care and Use Committee.

C57BL/6J mice were obtained as breeder pairs from the Jackson Laboratory (Bar Harbor, ME) and reared at the University of Missouri. Approximately equal numbers of six-to-ten week old age- and sex-matched mice were used in each experiment. The MAR1-5A3 and IgG1 isotype control antibodies were obtained from Bio X Cell (Lebanon, NH) [43]. Antibodies were intraperitoneally injected into mice at a dose of 2.5mg/mouse approximately 18 hours before mice were infected. Mice were intranasally infected with a dose of 2.5x10^7^ CFU/30μL. All infected mice were monitored by daily assignment of health scores, which involved assessments of their appearance and activity. Animals that survived to the end of the 14-day observation period or were identified as moribund (defined by pronounced neurologic signs, inactivity, and severe weakness) were euthanized by CO_2_ asphyxiation followed by bilateral pneumothorax or cervical dislocation, according to the American Veterinary Medical Association Guidelines on Euthanasia.

### Histology processing and scoring

After intranasal infection of *Yersinia pestis*, mice were euthanized, tissues were removed and fixed in 10% zinc formalin for a minimum of 24 hours, then sectioned and stained with hematoxylin and eosin (H&E). For severity scoring, H&E-stained slides were blinded. For the liver, three low-power fields were quantified and averaged to obtain a lesion severity score: inflammatory foci and necrotic foci were evaluated by size and frequency in each tissue section and each was assigned a score from 0-5 (1: small foci, only 1 or 2 in the section, 5: many large foci throughout the section) then added together for a final score. For the spleen, necrotic tissue and inflammatory foci with or without increased megakaryocytes were observed and evaluated for the degree to which they were present (0-5 scale).

### Serum Analyses

Cytokines: Tissues were homogenized in 1mL of PBS, whereas serum was obtained after clotting of red blood cells. Cytokines (IL6, TNFα and IL10) were quantified by ELISA kit (R&D Systems, MN) per manufacturer instructions. For analysis of serum chemistry, on day 5 post-infection, blood was collected in heparin tubes and centrifuged for 2 minutes at 14,000 rpm to separate serum, then stored at -80°C until analysis.

### Statistical Analysis

Data from all trials were used in statistical evaluation of data. Survival data were routinely analyzed by Gehan Breslow log rank test; bacterial load, cytokine, and serum enzyme data were not normally distributed and were analyzed by Mann Whitney test; histopathology was analyzed by Student’s t test. Significance was concluded when P<0.05.

## Results

### Antibody neutralization of IFNAR protects mice from plague

To investigate the potential of therapeutic targeting of IFNAR as a plague treatment, we treated wild type C57BL/6 mice with MAR1, a monoclonal neutralizing antibody against murine IFNAR or with non-specific IgG1 as an isotype control [43]. Mice were treated 18 hours prior to intranasal challenge with *Y. pestis* KIM5-, then monitored for disease over a 14-day period. In this model, the non-pigmented mutant *Y. pestis* causes only minor infection in the lungs and spreads systemically to cause lethal secondary septicemic plague [44]. As anticipated, a single treatment of MAR1 prior to infection was sufficient to protect mice from developing lethal disease, with a significant increase in survival compared to the IgG isotype-treated control mice (Figure 1A). We measured bacterial titer from the lungs, liver and spleen of these mice. The impact of MAR1 treatment on bacterial growth appeared to be a small improvement in clearance in the lungs and spleen, though the P-value was 0.1 for both tissues between treatment groups (Figure 1B). For the liver, however, we found higher numbers of MAR1-treated mice that had cleared the infection, but the median bacterial load recovered from the infected livers was higher than the IgG-treatment group suggesting a biphasic response. We also analyzed inflammatory and anti-inflammatory cytokines in the serum and spleen on day 5 post-infection. The levels of pro-inflammatory IL6 were high in both treatment groups for the serum and spleen (Figure 1C). In striking contrast, serum TNFα was markedly reduced by MAR1 treatment whereas splenic TNFα was not impacted (Figure 1D). Like IL6, the anti-inflammatory IL10 was also not impacted by MAR1 treatment in the serum or spleen (Figure 1E). Combined, the data suggest that early MAR1 treatment may improve the anti-bacterial response in the lungs and spleen while having a major impact on reducing serum TNFα, which correlated with improved survival.

**Figure 1.**
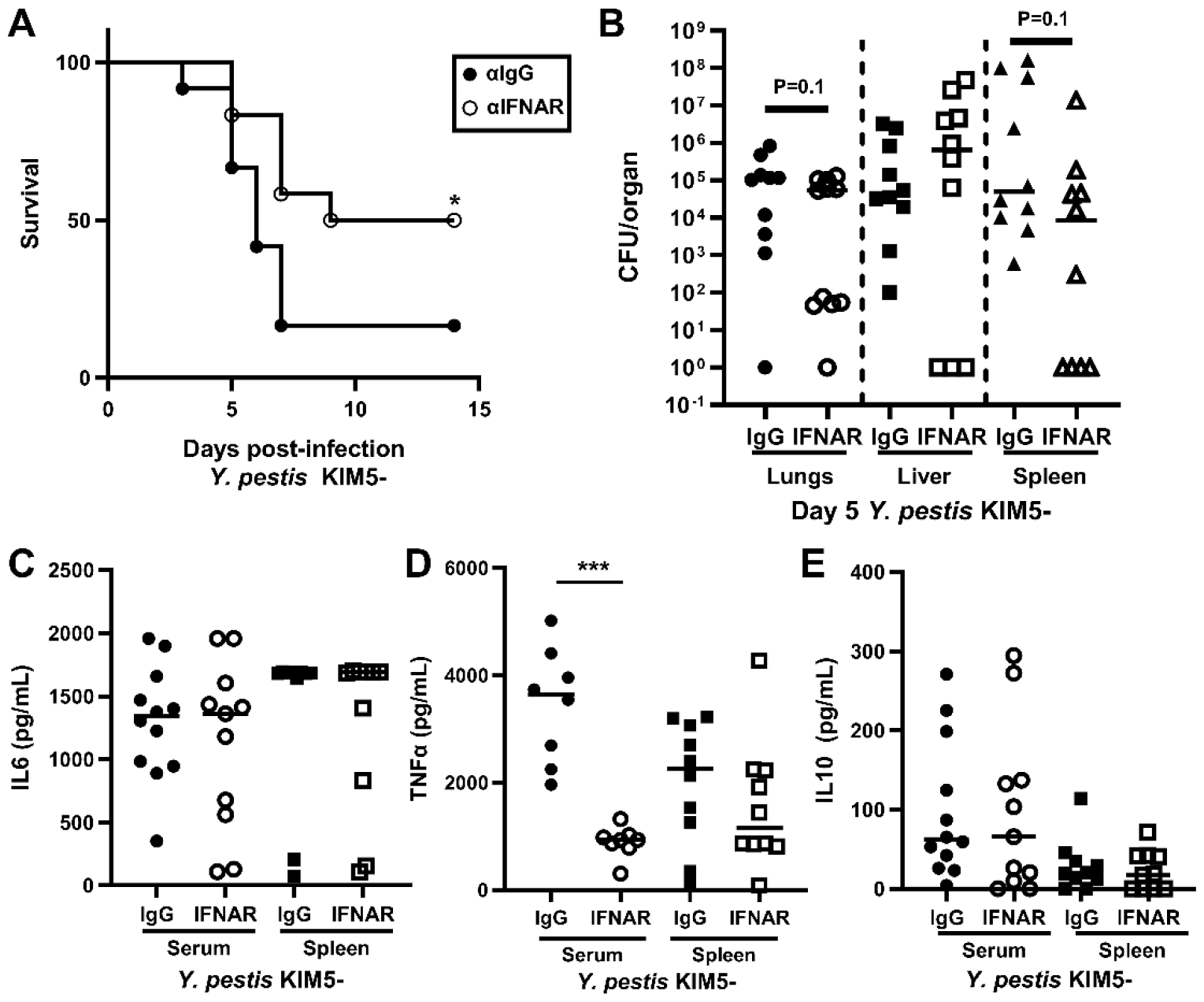
Antibody inhibition of IFNAR is protective in a murine pulmonary infection model of plague by reducing serum TNF_α_. Groups of 3-6 wild type C57BL/6 mice were treated with monoclonal IFNAR antibody MAR1 (open symbols) or anti-IgG1 (closed symbols) 1 day before intranasal challenge with *Y. pestis* KIM5-. A) Mice were monitored for development of plague over 14 days; n=12 per group, collected in 3 independent trials. B) Bacterial titers in lungs (circles), livers (squares) and spleens (triangles) were determined on day 5 post-infection. (C-E) Serum (circle) and spleen (square) cytokine levels on day 5 post-infection: C) IL6, D) TNFα, E) IL10. Data shown were collected in 2-3 independent trials, n=10-12 per group. Data were evaluated by Gehan Breslow log rank test (A) or by Mann-Whitney (B-E); *P<0.05, ***P<0.001.

### Y. pestis YopJ is required for the pathologic role of type I IFN

To determine if YopJ was important to the pathologic role of IFNAR, we repeated this study using *Y. pestis yopJ*. Mice were treated with IgG or MAR1, then infected with *Y. pestis yopJ* and observed for 14 days for development of disease. Comparing the survival of control mice challenged by *Y. pestis yopJ* to mice challenged with KIM5-, we found reduced mortality in the *yopJ*-infected group (Figure 2A). Though the reduced mortality was reproducible, this study showed a P-value of 0.1, consistent with a small attenuation of virulence caused by loss of YopJ activity. In striking contrast to what we observed for KIM5-infected mice, MAR1 treatment of *yopJ*-infected mice provided no detectable protection from mortality compared to IgG treatment (Figure 2B). Like the KIM5-infected mice, however, MAR1 treatment reduced bacterial load in the lungs of *yopJ*-infected mice (Figure 2C). Unlike the KIM5-infected mice, however, MAR1 treatment did not decrease bacterial load in the spleen, and if anything, there was increased *Y. pestis yopJ* in the spleen of MAR1-treated mice compared to IgG treatment. Bacterial load recovered from the liver seemed unaffected by MAR1 treatment. Similar to KIM5-infected mice, IL6 levels in the serum or spleen of *yopJ*-infected mice were not impacted by MAR1 treatment (Figure 2D). Somewhat unexpectedly given the lack of protection, however, there was significantly reduced serum TNFα in MAR1-treated *yopJ*-infected mice compared to αIgG treatment whereas splenic TNFα was similar between the treatment groups (Figure 2E). In contrast, no detectable differences were observed in IL10 in the serum or spleen between treatment groups (Figure 2F). Combined, these data suggest that IFNAR contributes to serum TNFα regardless of the presence of YopJ, whereas IFNAR contributes to bacterial clearance in the spleen only when YopJ is absent. These observations suggest that YopJ blocks an anti-bacterial role for the type I IFN response, whereas the impact of this response on the disease-phase cytokine storm may be independent of YopJ. These two opposing host responses may therefore underlie the apparent undetectable impact of IFNAR on survival of mice challenged with *Y. pestis yopJ*.

**Figure 2.**
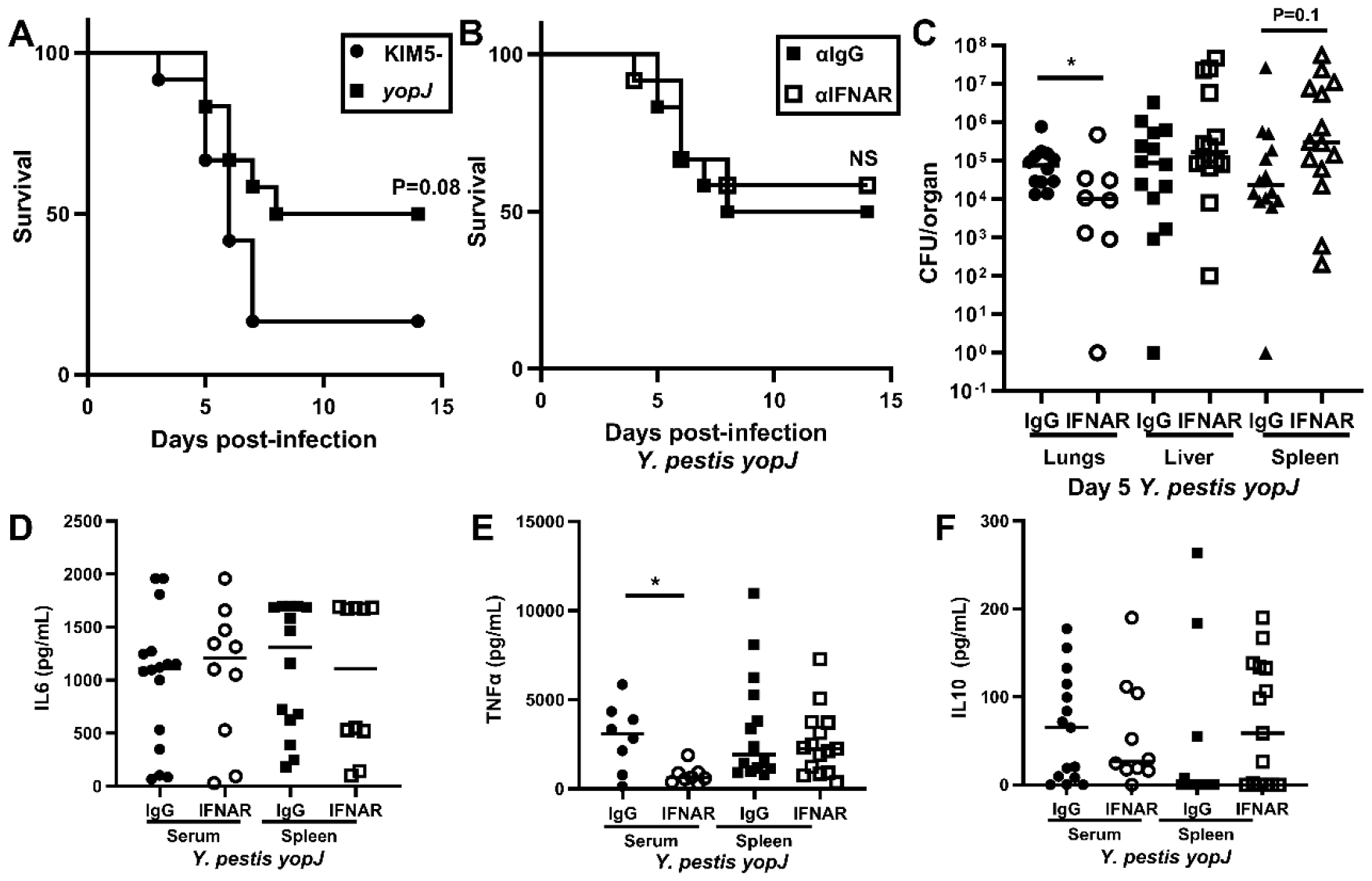
Antibody inhibition of IFNAR has no detectable impact on the virulence of Y. pestis YopJ_C172A_ in the mouse pulmonary infection model. Groups of 3-4 mice were treated with MAR1 (open symbols) vs IgG (closed symbols) antibodies 1 day before *yopJ*_*C172A*_ infection. A-B) Survival curves of A) KIM5- and *yopJ* infections in IgG-treated mice, and B) MAR1 and IgG treated mice infected with *Y. pestis yopJ*; data shown were collected in 3 independent trials, n=12 per group. C) Bacterial titer in the lung (circle), liver (square), and spleen (triangle) on day 5 post-infection; n=8-14 per group, collected in 2-3 independent trials; bars indicate median. (D-F) Cytokine levels in the serum (circle) and spleen (square) on day 5 post-infection: D) IL6, E) TNFα, F) IL10; data shown were collected in 2-3 independent trials, n=10-16 per group; bars indicate median. Data from A-B were evaluated by Gehan Breslow log rank test; data from C-F were evaluated by Mann-Whitney, *P<0.05, all other comparisons were not significant.

### IFNAR and YopJ cooperate in enhancing disease

Previous work identified a potential role for IFNAR signaling on the development of neutrophilic inflammatory foci in the *Y. pestis*-infected liver [39]. We therefore sought to understand if YopJ was also important for this phenotype. To address this, we used histochemistry on liver and spleen collected from infected mice on day 5 post-infection. In this assessment, we commonly observed severe necrosis in the spleens of MAR1 and IgG-treated groups infected with KIM5-(Figure 3A-B). Overall it appeared that the absence of *yopJ* resulted in improvement in the severity scores of spleens, with the lowest median score in the IgG/*yopJ* group, though there were no significant differences detected by any comparison (Figure 3C, E). Inhibition of IFNAR had an apparent negative effect on pathology in the *yopJ*-infected mice (Figure 3D-E). This data is consistent with YopJ-interference with the type I IFN response in the spleen and suggest that YopJ-inhibition of IFNAR in the spleen promotes virulence.

**Figure 3.**
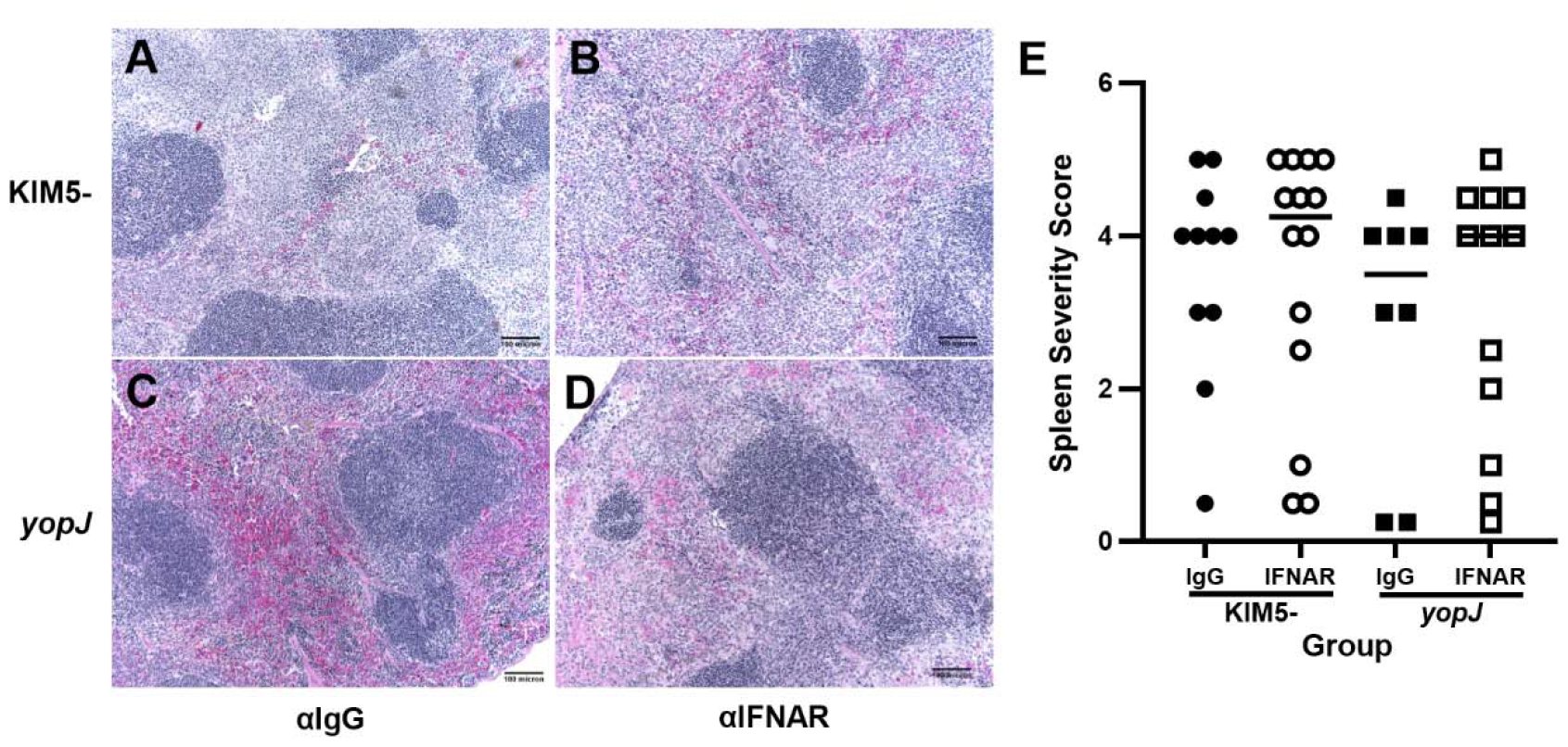
Inhibition of IFNAR has minimal impact on pathology in the spleen. Groups of 3-4 mice were treated with MAR1 (B, D, open symbols) or IgG (A,C, closed symbols) antibodies 1 day before intranasal infection with *Y. pestis* KIM5-(A,B, circles) or the *yopJ* mutant (C-D, squares). On day 5 post-infection, mice were euthanized, tissues removed and processed for histopathology analysis. (A-D) Representative images of H&E-stained spleens of mice; (E) Samples were blinded and scored for lesion severity. Data shown were pooled from 4 independent trials, n=7-14 per group. (E) Data were evaluated by Student’s t test, comparing IgG to MAR1 for each bacterial strain, no significance was detected between any groups.

Liver tissue sections were also scored for lesion severity to include necrotic as well as inflammatory foci, factoring in the size and number of each. With this analysis, we found that YopJ activity leads to reduced lesion severity in the liver, with increased median severity score in IgG-or MAR1 treated *yopJ*-infected mice compared to the KIM5-infected groups (Figure 4A-E). This data is consistent with the ability of YopJ to suppress inflammation. However, these results do not correlate with the survival data, suggesting that the inflammatory and necrotic lesions in the liver do not cause the survival differences observed on MAR1 treatment.

**Figure 4.**
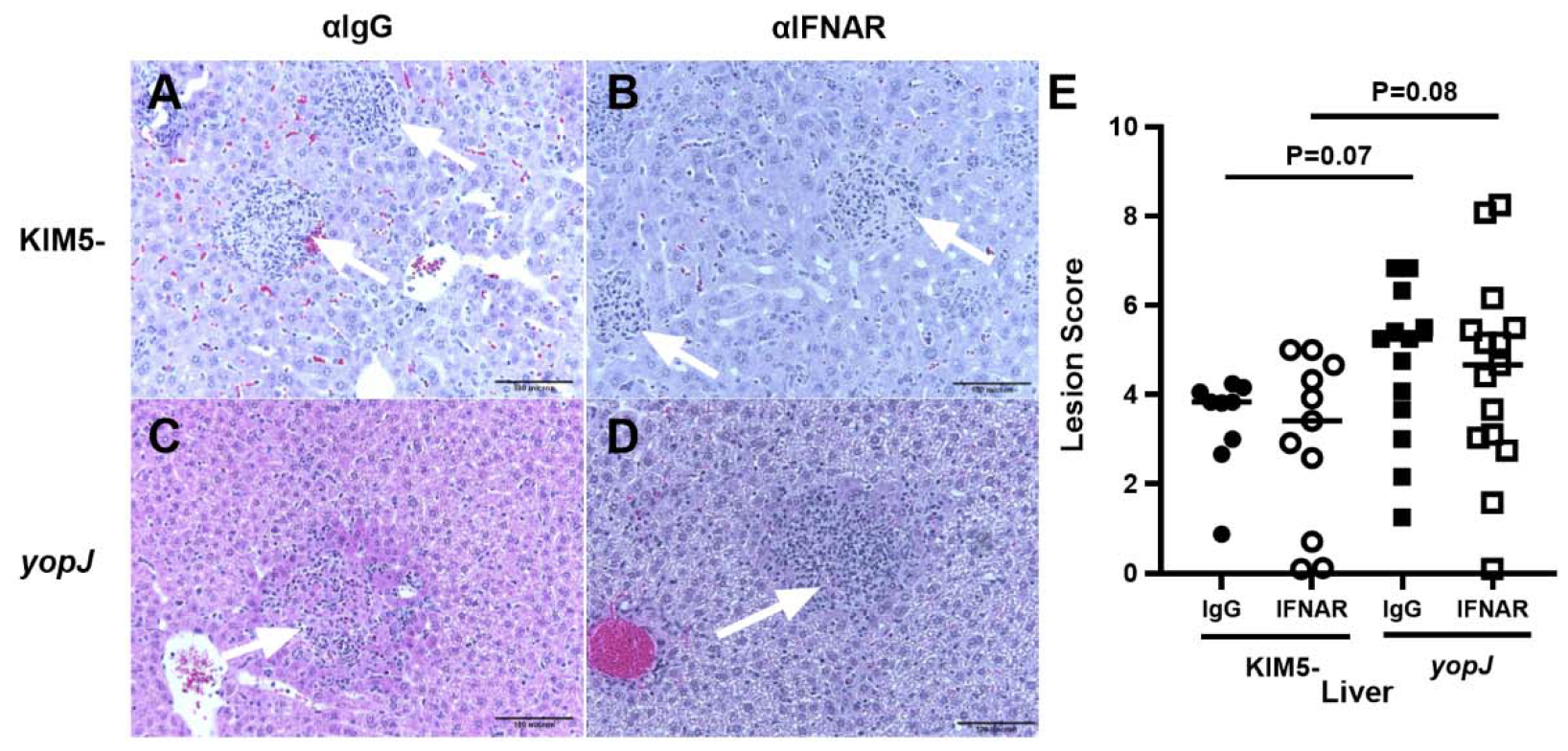
YopJ but not IFNAR impacts inflammation in the liver. Mice were treated with MAR1 (B, D, open symbols) or IgG (A,C, closed symbols) antibodies 1 day before intranasal infection with *Y. pestis* KIM5-(A,B, circles) or the *yopJ* mutant (C-D, squares). On day 5 post-infection, mice were euthanized, tissues removed and processed for histopathology analysis. (A-D) Representative images of H&E-stained livers of infected mice; (E) Samples were blinded and scored for lesion severity, including inflammatory foci (size and frequency, shown by white arrows) and hepatocyte necrosis. Data shown were pooled from 5 trials, n=9-15 per group. (E) Data were evaluated by Student’s t test, comparing IgG to MAR1 for each bacterial strain.

To assess potential loss of liver function, we analyzed day 5 serum for the following parameters: Glucose, urea nitrogen (BUN), albumin, total protein, alanine aminotransferase (ALT), aspartate aminotransferase (AST), bilirubin, and alkaline phosphatase (ALP). Of these, all infected groups showed significant up-or down-regulation of glucose, total protein, albumin, AST, ALT, and ALP compared to the control groups that were not infected (Table 1). In contrast, bilirubin was not detectably different between any group. Urea nitrogen was significantly decreased in the MAR1 treatment group compared to not-infected whereas the average BUN values for the IgG-treated, KIM5 infected group was not significant compared to uninfected. Overall, the serum chemistry suggested that mice in all 3 groups were experiencing acute phase responses with potentially decreased liver function, with the MAR1-treated, KIM5-infected having the least severe disease.

**Table 1.** Serum chemistry analysis of anti-IFNAR treatment on Y. pestis KIM5- and yopJ infection.

|  | Not infected | IgG:KIM5- | MAR1:KIM5- | IgG:YopJ |
| --- | --- | --- | --- | --- |
| Glucose (mg/dL) | 363 | 79**** | 115**** | 102**** |
| Urea Nitrogen (mg/dL) | 32 | 27 | 20* | 25 |
| Albumin (g/dL) | 3.2 | 2.7** | 2.8* | 2.9 |
| Total Protein (g/dL) | 5.3 | 6.4** | 6.6** | 6.5** |
| AST (U/L) | 133 | 730*** | 556* | 521<sup>‡</sup> |
| Bilirubin (mg/dL) | 0.3 | 0.5 | 0.3 | 0.5 |
| ALT (U/L) | 36 | 166* | 138<sup>‡</sup> | 189* |
| ALP (U/L) | 101 | 24**** | 33**** | 28**** |

Statistical Evaluation by one way ANOVA followed by Tukey's; \*P<0.05, \*\*P<0.01, \*\*\*P<0.001, \*\*\*\*P<0.0001, <sup>‡</sup>P=0.1, compared to uninfected.

## Discussion

The goal of this study was to examine therapeutic inhibition of type I IFN signaling and its impact on host-pathogen interactions that contribute to the progression of plague. We established for the first time that a single dose of neutralizing IFNAR antibody is sufficient to protect mice from developing lethal plague. Given that the half-life of the MAR1 IgG1 antibody is estimated to be 6-8 days, it is difficult to ascertain the kinetics whereby this treatment is therapeutic from this study. We found that following this single dose administration of MAR1, pleiotropic effects were observed in infected tissues. Despite an impact on bacterial growth in the lungs and spleen, however, we did not detect large histopathological indicators that might implicate neutrophil recruitment as an underlying mechanism of protection. Rather, the systemic effects of the type I IFN response may have been driven by TNFα, as substantial reduction of serum TNFα was observed in the MAR1-treated mice. This is somewhat surprising, however, because early administration of TNFα was shown to be protective in a mouse septicemic plague model [45]. This apparent contradiction in data may be explained by tissue-specific responses that may be missed when *Y. pestis* is directly inoculated in blood as it was done in the previous study. Alternatively, TNFα may induce anti-bacterial effects when present early in the infection as it was previously administered, before the animals enter the disease phase, whereas systemic TNFα during the disease phase may worsen the progression of sepsis.

Type I IFNs regulate autocrine and paracrine responses that influence disease outcome [46]. Although TNFα may not be directly controlled as an ISG, there are numerous studies describing co-dependencies between the IFN response and the TNFα response. For example, dendritic cells, macrophages and likely other cell types produce IFN-dependent TNFα which has been associated with worsening disease or pathology [47-49]. Because *Y. pestis* targets phagocytic cells for injection of Yops and we previously identified an IFN-dependent reduction in the neutrophil population in the bone marrow, we think it is likely that the pathologic type I IFN response impacts neutrophil recruitment. This is supported by reduced bacterial clearance observed in the lungs and spleen of IFNAR-treated mice. However, since the non-pigmented *Y. pestis* KIM5-strain that was used in this study is not able to cause disease in these mice, the IFNAR-dependent clearance may not impact outcome [44].

There are likely additional systemic effects of the type I IFN response that were not measured here that could negatively affect the severity of sepsis. For example, recent work has established a linkage between IFN signaling and coagulation and thrombosis. Following cecal ligation and puncture in the mouse model, an excessive or dysregulated type I IFN response activated pyroptosis in macrophages causing extracellular vesicle (EV) release which was demonstrated to increase circulation of coagulation factors [50]. The resulting thrombosis induced the progression of disseminated intravascular coagulation (DIC). This mechanism may be relevant to *Yersinia* infection, as the T3SS is known to activate the inflammasome, and YopJ is known to cause pyroptosis. Perhaps IFNAR and YopJ coordinate in this manner to further the progression of DIC.

Overall it appears that tissue specific and systemic effects of the type I IFN response influence *Y. pestis* infection in the mouse model. Future work to examine cell-type specific responses will be needed to separate the pathologic from beneficial type I IFN responses, as it appears that therapeutic targeting of one or more of these responses may improve the outcome of plague.

## Acknowledgements

We are grateful to Dr. Craig Franklin for advice on histopathology, to Dr. Carol Reinero for advice on serum chemistry, and to members of our laboratory for helpful discussions. H&E-stained slides were prepared by IDEXX-RADIL (Columbia, MO); serum chemistry was conducted at the MU Veterinary Medical Diagnostic Laboratory (Columbia, MO).

